# Genetic determinants of host tropism in *Klebsiella* phages

**DOI:** 10.1101/2022.06.01.494021

**Authors:** Beatriz Beamud, Neris García-González, Mar Gómez-Ortega, Fernando González-Candelas, Pilar Domingo-Calap, Rafael Sanjuan

## Abstract

Bacteriophages play key roles in bacterial ecology and evolution and are potential antimicrobials. However, the determinants of phage-host specificity remain elusive. Here, we used 46 newly-isolated phages to challenge 138 representative clinical isolates of *Klebsiella pneumoniae*, a widespread opportunistic pathogen. Spot tests revealed a narrow host range for most phages, with <2% of 6319 phage-host combinations tested yielding detectable interactions. Bacterial capsule diversity was the main factor restricting phage host range. Consequently, phage-encoded depolymerases were key determinants of host tropism, and we identified depolymerase sequence types associated with the ability to infect specific capsular types across phage families. Phages showing a capsule-independent mode of entry exhibited a much broader host range, but their infectivity was still restricted by complex intracellular defense mechanisms. These findings expand our knowledge of the complex interactions between bacteria and their viruses, and have implications for the biomedical and biotechnological use of phages.

## INTRODUCTION

Bacteriophages are extremely abundant and diverse biological entities and, as such, they are key ecosystem actors (Abedon, 2008). Yet, the mechanisms that govern phage-bacteria interactions are not well understood, since most studies have focused on small sets of virus-host pairs or have not followed a systematic whole-genome approach (Chevallereau et al., 2022; Lamy-Besnier et al., 2021). Advances in high-throughput sequencing and metagenomics have allowed linking phages to their hosts via genomic signatures (Edwards et al., 2016; Roux et al., 2015). Although very useful, these methods do not provide individual information about phage-bacteria interactions. Thus, isolation of phages using strains of interest (Maffei et al., 2021; Porter et al., 2020) along with genome screenings (Mutalik et al., 2020) can complement these studies to better understand phage-bacteria interactions.

The determinants of phages’ host range are complex. Typically, a phage infects only a subset of strains of a given bacterial species (de Jonge et al., 2019). However, some phages can infect different bacterial species (Göller et al., 2021; Kauffman et al., 2022) or even distinct genera (Gambino et al., 2020; Hamdi et al., 2017). Available tools can predict phages’ host range with moderately good accuracies at the family and genus levels, but perform worse at lower taxonomical ranks (Coutinho et al., 2021; Villarroel et al., 2016; Young et al., 2020). Bacteria deploy a variety of mechanisms that can restrict phage infection, ranging from modification of surface receptors to a continuously expanding number of immunity systems, such as regularly-interspaced short palindromic repeats (CRISPRs), restriction-modification (RM) systems, abortive infection systems, and prophage super-immunity, among many others (Bernheim and Sorek, 2020; Hampton et al., 2020), complicating our understanding of phage-host interactions.

Bacterial capsules are another complex determinant of phage tropism, since they can protect bacteria from infection by masking receptors (Scholl et al., 2005) or being used for phage attachment (Porter et al., 2020). Most environmental and opportunistic bacteria present an extracellular capsule (Rendueles et al., 2017). Capsules protect the cell from the immune system of multicellular organisms being important virulence factors (Mostowy and Holt, 2018). To overcome the bacterial capsule, some phages encode depolymerases capable of digesting specific oligosaccharide bonds (Latka et al., 2019; Pires et al., 2016). Most phages contain one depolymerase and, as a result, infect one or a few cross-reactive capsular types (Pieroni et al., 1994). Depolymerases have a modular structure with potential module swapping between phages through recombination (Latka et al., 2021; Scholl et al., 2002). However, how binding and depolymerization determine successful phage infection is not well understood (Born et al., 2014; Pelkonen et al., 1992).

*Klebsiella pneumoniae* is a ubiquitous opportunistic Gram-negative *Enterobacteriaceae*, and is included in the ESKAPE pathogen group. *K. pneumoniae* offers an excellent model for studying the role of bacterial capsules in phage infectivity because it exhibits a remarkable capsular diversity, with 77 serotypes and over 180 capsular locus types (CLTs) described so far (Lam et al., 2022), in addition to a large repertoire of other phage defense mechanisms (Tesson et al., 2022). Given the limited treatments for emerging multidrug-resistant strains, *K. pneumoniae* is a major target of ongoing research on phage therapy (Eskenazi et al., 2022; Townsend et al., 2021; Herridge et al., 2020). Most *Klebsiella* phages infect a few CLTs, but their association with phage depolymerases sequences remains elusive (de Sousa et al., 2020; Venturini et al., 2020) or is restricted to a few phages (Domingo-Calap et al., 2020; Sørensen et al., 2021). Instead, the typical procedure involves extensive host-range testing along with the expression and purification of the candidate enzymes (Pan et al., 2017). Additionally, some phages without depolymerases have been described (Majkowska-Skrobek et al., 2021).

Here, we used a comprehensive collection of sequenced clinical *K. pneumoniae* strains (n = 138) and diverse novel environmental phages (n = 46) to study the relative importance of different bacteria and phage genetic traits as determinants of phages’ host range and infection outcome. We tested all phage-bacterium pairs by spot-testing and confirmed positive results by bactericidal, progeny production, capsule depolymerization, and adsorption assays.

## RESULTS

### Host tropism of isolated *Klebsiella* phages

We used water and soil samples near a sewage plant in Valencia (Spain) to isolate 70 plaques on 36 different *K. pneumoniae* clinical strains. Phages were amplified, purified, and sequenced, yielding 46 distinct genomes (**Figure S1** and **Table S1**). Phages belonged to 13 groups according to pairwise intergenomic similarity (IGS >= 45%). We named each virus with the first letter of the associated viral family, the number of the phylogenetic group, and a letter to identify each species (IGS >= 95%) (**Figure 1**). All phages lacked integrases, although several had lysogeny marker genes such as transcriptional repressors or ParA/B genes (**Figure 1** and **Table S1**). Transmission electron microscopy revealed different morphologies of representative phages (**Figure S2**). To determine their taxonomic classification, we retrieved 478 phage genomes from databases exhibiting at least 30% coverage and 70% average nucleotide identity (ANI) with our phages. According to pairwise IGS values, the 46 phages represented five families of the *Caudovirales* order (*Autographiviridae, Drexlerviridae, Myoviridae, Siphoviridae*, and *Podovirida*e) and a variety of genera *(Drulisvirus, Przondovirus, Webervirus, Gamaleyavirus, Mydovirus, Yonseivirus, Jedunavirus*, and *Roufvirus;* **Figure 1** and **Table S1**). They encompassed most families and subfamilies of previously described *Klebsiella* phages (**Figure S3**). All the phages except one exhibited < 95% IGS with available sequences, suggesting that they represented novel species. Some phages were unclassified at lower taxonomic ranks, which led us to suggest that they represented three new subfamilies and five new genera (**Table S1** and **Figure S4**).

**Figure 1.**
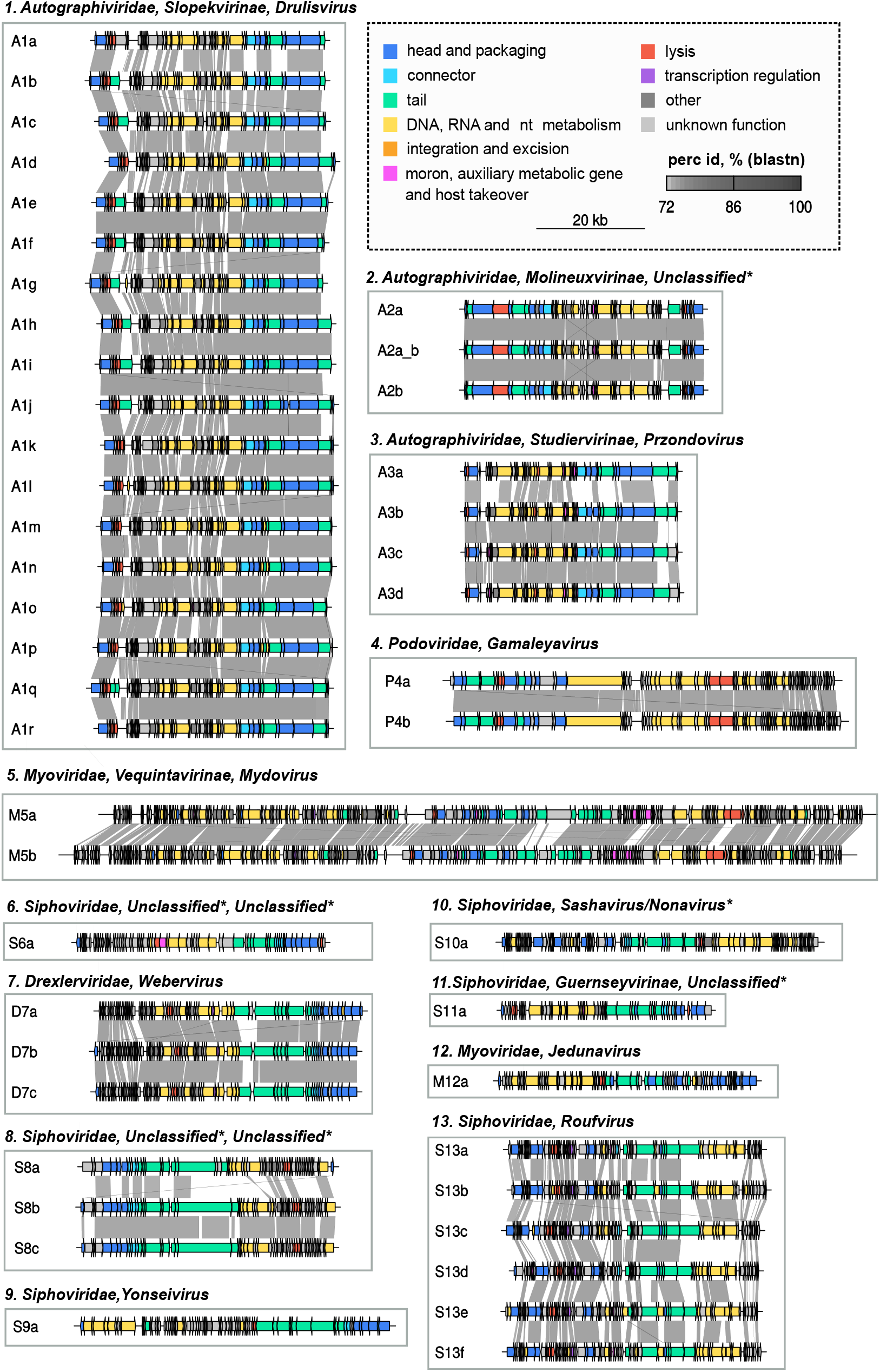
Diversity of isolated *Klebsiella* phages. IGS values were used to define 13 similarity groups and to classify phages in the closest viral family, subfamily (if applicable) and genus after comparison with database sequences. Phages were named after the first letter of the associated viral family, the number of the phylogenetic group and a letter to identify each species or strain. A: *Autographiviridae*; D: *Drexlerviridae*, M: *Myoviridae*; *P: Podovirida*e; S: *Siphoviridae*. The genome organization of each phage is shown. Arrows represent coding sequences (CDSs) and are colored based on functional categories of PHROGS.

To examine the host range of isolated phages, we selected 138 sequenced *K. pneumoniae* clinical isolates from the Valencia region of 59 different CLTs, 14 O locus types (OLTs), and 76 sequence types (STs; **Table S2**). Of them, 16 were used for the initial isolation of phages, along with 20 additional strains not included in the collection (**Table S2**). The 138 strains were diverse in terms of antibiotic susceptibility, presence of prophages, and anti-phage defenses such as CRISPR-Cas and restriction-modification systems (**Figure S5**). The resulting collection represented the overall species’ diversity, as shown by comparison with the main clonal groups of *K. pneumoniae* (**Figure S6**).

To determine host-tropism, we performed triplicate spot tests for all the 46 × 138 = 6348 phage-bacteria pairs (6319 after removing the phage-bacteria combinations used for initial phage isolation to avoid sampling bias). Only 124 combinations yielded reproducible spots, suggesting that, on average, a *Klebsiella* phage can infect less than 2% of host strains (**Figure 2** and **Table S3**). Host breadth ranged from 0 to 19 of the 138 strains (excluding the isolation strain), with two distinct patterns: most phages (42/46) infected one or a few strains (median: 1, mean: 1.79; 1-3 CLTs), but the four phages in groups 8 and 9 (S8/S9 phages) exhibited a much broader host range (median: 11.5, mean: 12.25; 4-16 CLTs, **Figure 2** and **Table S3**).

**Figure 2.**
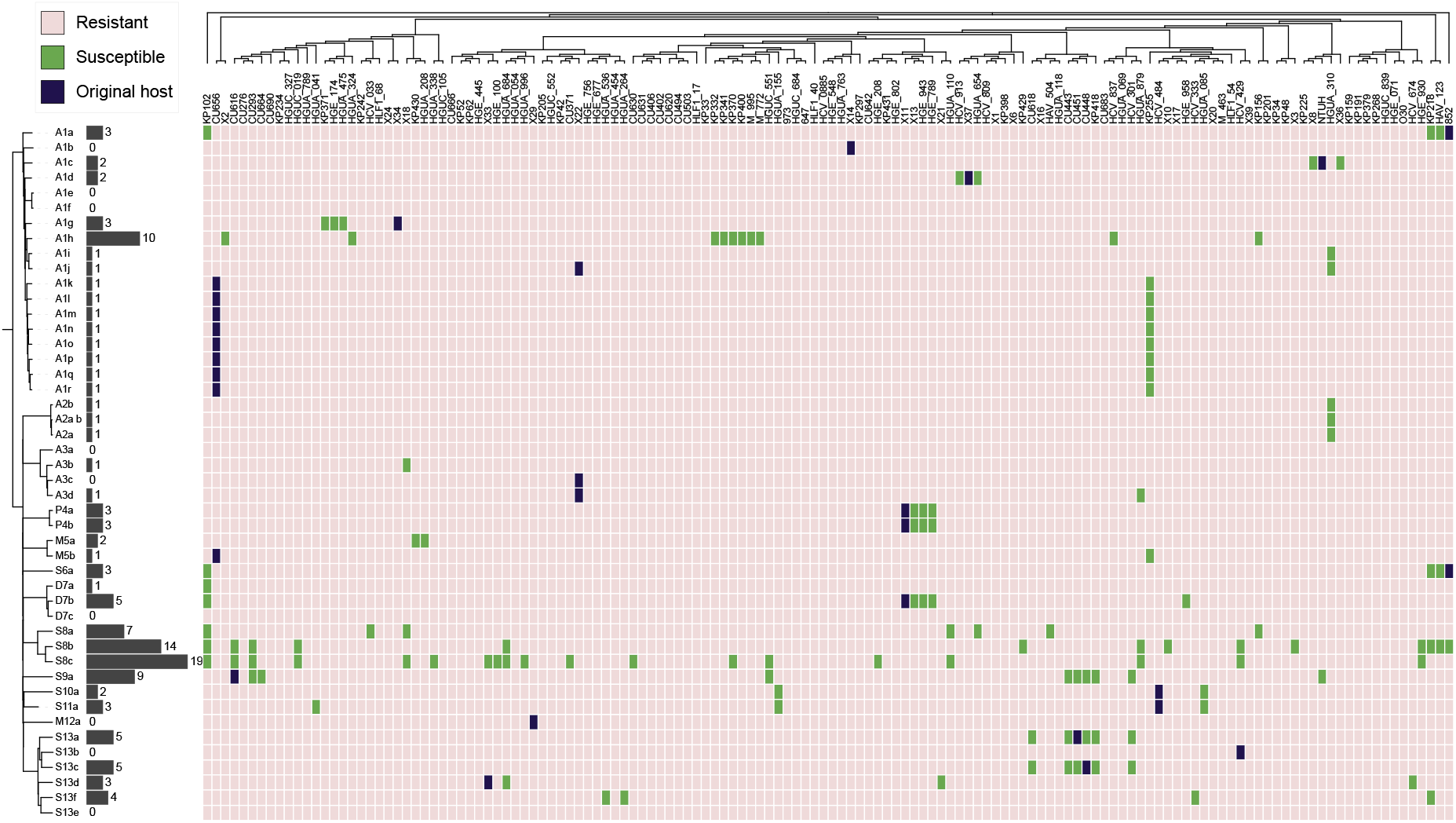
Phage host tropism towards *K. pneumoniae* collection. Each row represents a phage (n=46) and each column a bacterial strain (n=138). For phages, the dendrogram constructed from IGS values is shown, whereas the ML core phylogeny is shown for bacteria. A given bacterial strain was considered to be susceptible to a given phage (green boxes) if at least two thirds replicates of the spot assay were positive. Original host-phage pairs in which each phage was primarily isolated are indicated in purple. The black histogram and associated numbers indicate the host breadth of each phage in the 138 total bacterial strains analyzed excluding isolation hosts.

### Depolymerase-capsule interactions are the main determinants of host tropism

To investigate the determinants of host tropism, we first focused on the 42 non-S8/S9 phages. We assessed which host features predicted spotting by phages (true positive rate, TPR). For each trait, we sampled random reference hosts and the phages’ spot outcomes and used this information to predict the ability of phages to spot other bacterial strains sharing the same trait. This revealed that CLT was the host trait that best predicted tropism. Specifically, a phage spotting in a given strain had a 92% probability of also spotting in other strains of the same CLT (TPR = 0.92 ± 0.001 for non-S8/S9 phages; **Figure 3A**). Bacterial ST was another good predictor of host tropism (TPR = 0.60 ± 0.01), albeit it is correlated to CLT in our dataset (Cramér’s V ES = 0.9). In contrast, OLT and 45 additional proteins encoding putative phage receptors were poor predictors of host tropism (TPR = 0.04-0.09). This confirmed that the bacterial capsule is a major determinant of host tropism in *Klebsiella* phages.

**Figure 3.**
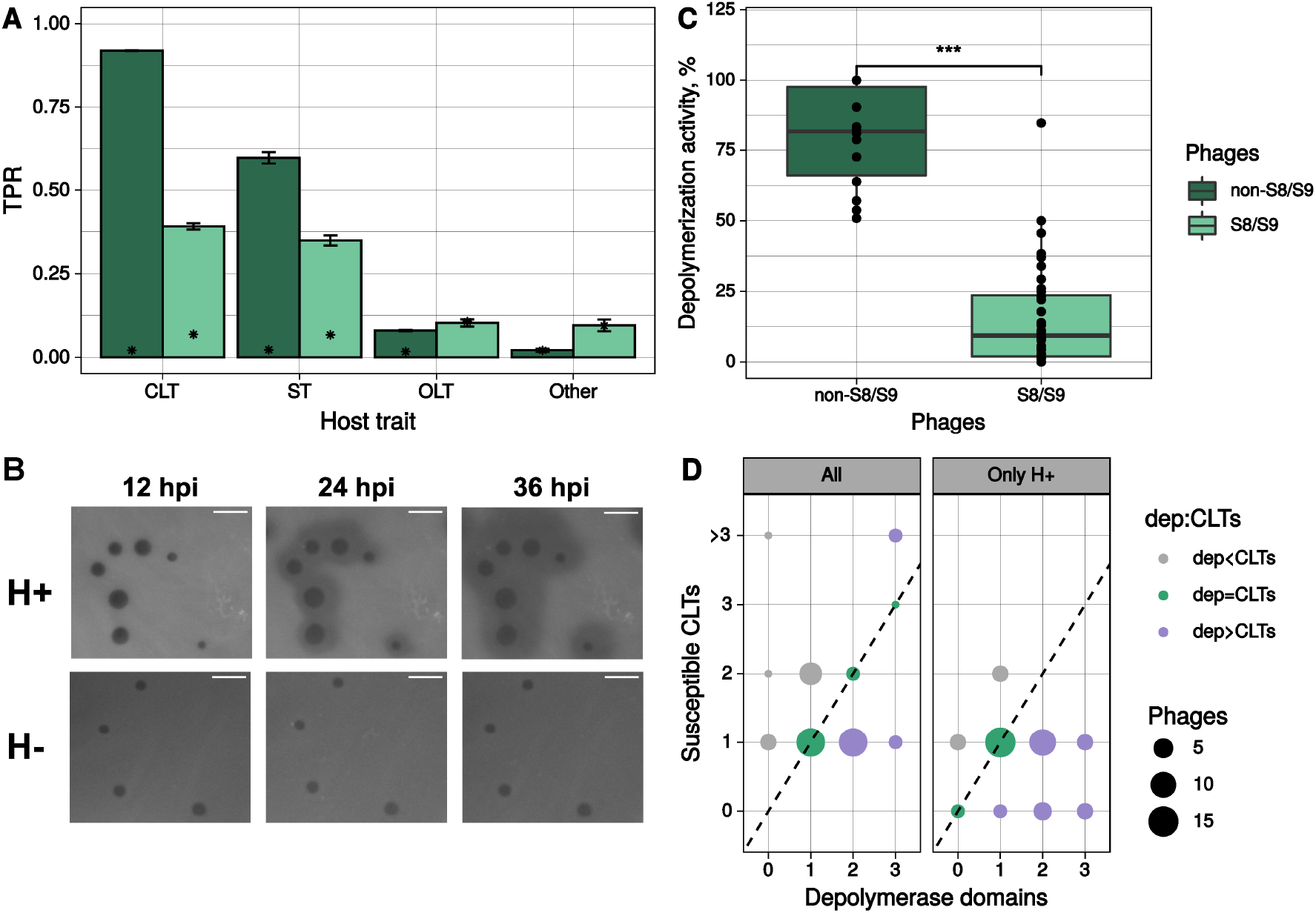
Depolymerase-capsule interactions are the main determinant of host tropism. **A**. Sensitivity or True Positive Rate (TPR) of different host traits to predict positive spot tests for non-S8/S9 phages and phages from groups S8/S9. Bars represent the standard error of 3 independent calculations. Asterisks represent the mean TPR values obtained with randomized data (null expectation). CLT: capsular locus type; OLT: O-antigen locus type; ST: sequence type. Other: presence of 45 secondary receptors (pooled). **B**. Example of phage plaques surrounded by haloes that keep expanding with increased incubation times (H+) or not (H-). hpi: hours post-infection. Scale bar: 5 mm. **C**. Depolymerization activity of halo-producing non-S8/S9 phages and non-halo-producing S8/S9 phages relative to non-inoculated controls. Each point represents the mean activity of a given phage-bacterium combination tested. Asterisks indicate statistical significance (t-test: P < 0.001). **D**. Susceptible CLTs by phage depolymerase dosage considering all CLTs or those CLTs in which phages produced a halo (H+). Points with >3 CLTs only included S8/S9 phages. The dashed lines show the expected 1:1 relationship between depolymerase domains and susceptible CLTs.

Phages infecting encapsulated bacteria can encode depolymerases that digest capsular polysaccharides to provide access to secondary receptors on the cell surface (Latka et al., 2019). We found depolymerase domains in 38 of the 42 non-S8/S9 phage genomes. These were usually in the same ORFs as tail fiber/spike proteins. Many phages (20/42) encoded a single putative depolymerase, but 15 phages encoded two and 3 phages encoded three (**Table S1**). Most of these depolymerases should be functional because plaques were typically surrounded by haloes (**Table S1** and **Figure 3B**), a typical indicator of depolymerase activity. Supporting this, quantitation of capsule polysaccharide levels showed a strong reduction (t-test P < 0.001) in the 14 bacterial strains inoculated with halo-producing phages relative to non-inoculated controls (**Figure 3C**). Failure to detect depolymerases in some phages can be due to high sequence divergence or annotation problems (de Sousa et al., 2020; Townsend et al., 2021). For instance, three of the non-S8/S9 phages without detectable encoded depolymerases showed haloes and significant activity against the capsule, suggesting some depolymerase activity (**Figure 3C**). However, in most cases, the number of depolymerases encoded was equal to or larger than the number of susceptible CLTs, especially when considering CLTs in which the phage produced a halo (**Figure 3D** and **Table S1**), suggesting inactivity of some depolymerases or specificity of action towards CLTs not included in this work.

### Broad-range S8/S9 phages exhibit capsule-independent tropism

Whereas in our dataset, non-S8/S9 phages infecting a specific strain had a 92% probability of also infecting other strains of the same CLT, this association dropped for the broad-range S8/S9 phages (TPR = 0.39 ± 0.01; **Figure 3A**). Hence, the tropism of S8/S9 phages was weakly determined by CLT. Despite their broader host range, the S9a phage encodes no depolymerases and S8 phages encode three, similar to narrow-range phages from groups 1 and 7 (**Table S1** and **Figure 3D**). This suggests that S8/S9 phages use alternative mechanisms for penetrating the bacterial capsule. We also noticed that none of the S8/S9 phages produced plaque haloes (**Table S1** and **Figure 3D**). To explore this further, we experimentally analyzed the depolymerase activity of 36 different host-phage pairs involving S8/S9 phages. In contrast to the strong capsular polysaccharide digestion activity shown by the other phages, S8 phages exhibited variable depolymerization activity depending on the strain tested (**Figure 3C**), with variations found even among strains of the same capsular type. For instance, phage S8b depolymerized strain KL39-CU293 but did not show depolymerase activity in KL39-CU616. We found the opposite pattern in phage S8c, even though neither phage produced haloes in any of the tested hosts (**Table S1** and **Figure 3D**). Thus, some depolymerase domains found in the genomes of S8 phages should be active. However, in some hosts, these phages, as well as the S9a phage, may use alternative modes of entry that remain uncharacterized and presumably do not involve depolymerase activity.

### Sequence-based prediction of phage capsular tropism

The capsular type of bacteria can be obtained from their genome (Mostowy and Holt, 2018) but at present, it is not possible to predict the capsular tropism of phages based on its sequence. To address this, we first expanded our dataset by including 64 additional *Klebsiella* phages from the literature with known tropism. We then focused on the central and C-terminal regions of the tail-spike proteins where depolymerase specificity lies (Latka et al., 2019, 2021), what we referred to as receptor-binding domain (RBD). We extracted 59 RBDs from the 42 non-S8/S9 phages and 130 RBDs from additional *Klebsiella* phages. The 189 RBDs grouped into 39 clusters with at least two RBDs sharing > 40% sequence coverage and > 50% amino acid identity (**Table S4**). We found RBD clusters had a mean TPR of 53.0 ± 4.0% on phage tropism (**Figure 4A**, left panel). Remarkably, these values exceeded the TRP of phage phylogenetic markers. For instance, for any phage-CLT pair, other phages of the same phylogenetic group had only an 18% chance of infecting the same bacterial capsular type (**Figure 4A**, left panel). The accuracy of RBD-based CLT tropism prediction was variable, as 16 RBD clusters had a strong association with capsular tropism (TPR > 80%) whereas nine showed no association at all (**Figure 4A**, right panel). We expected this considering that (1) many phages contained several putative depolymerases, some of which could be cross-reactive against multiple capsular types, and (2) we could not experimentally test all the existing capsular types. Despite these limitations, we found 25 RBD clusters with a consensus capsular tropism, defined as clusters of RBD sequences in which at least two-thirds of the sequences belonged to phages spotting in the same CLT. These 25 clusters were composed of 79 RBDs of 65 phages and covered 19 distinct CLTs. RBD clusters involved phages from the same phylogenetic group (69.1%), but also from distinct genera and even families (22.7%; **Figure 4B**), revealing horizontal gene transfer of RBDs across large evolutionary scales.

**Figure 4.**
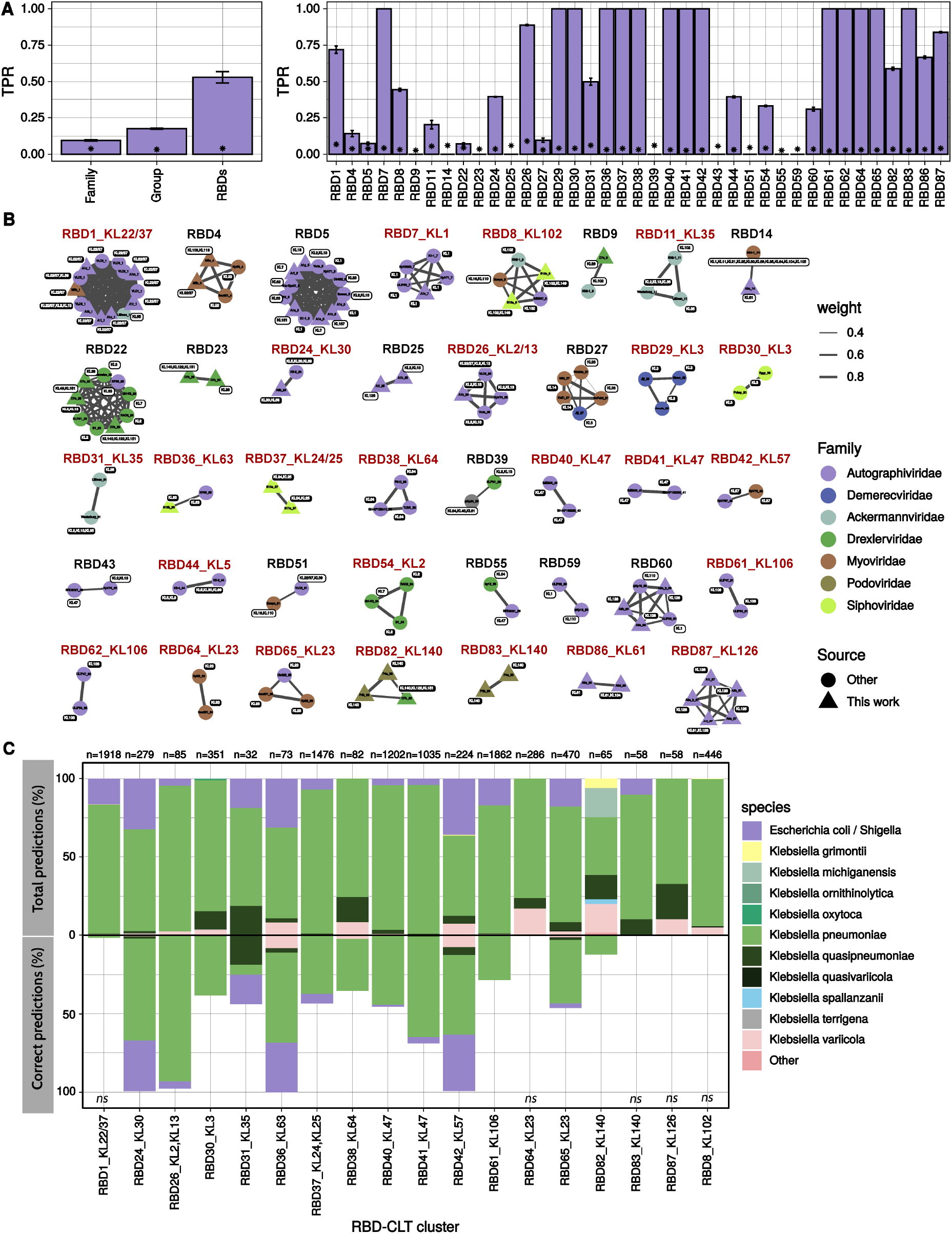
Sequence-based predictability of phage capsular tropism. **A**. Left panel: Sensitivity or True Positive Rate (TPR) of different phage traits to predict infections in capsular types of bacteria. Bars represent the standard error of 3 independent calculations. Asterisks represent the mean TPR values obtained with randomized data (null expectation). For RBDs, the average of the 39 RBD clusters is shown. Right panel: TPR for each individual RBD cluster. **B**. Representation of the 39 RBD similarity clusters obtained. RBDs are named by the phage name followed by the phage protein in which they were detected. Clusters labeled in red are those for which a consensus capsular tropism could be obtained. Colors of nodes indicate phage taxonomic families. Sequences obtained in the present work are shown as triangles and previous published sequences as circles. Edge weight (width) represents amino acid identity between RBDs within the same cluster. Each RBD is annotated with the corresponding phage CLT tropism. White labels represent tropisms that did not match the consensus CLT of an RBD cluster. **C**. Ability of the consensus RBD clusters to predict the CLT tropism of prophages obtained from the RefSeq database as described in the main text. The percentage of total and correct predictions by bacterial species is shown. The n-values indicate the total predictions for each RBD. “Other” refers to spp. within *Klebsiella* sp. and also from other genera. *ns* denotes no significance by Fisher’s exact test (P > 0.05).

To further analyze the predictive power of RBD clustering, we extracted a representative sequence from the 25 RBD clusters showing a consensus capsular tropism (**Figure 4B**). We then interrogated the RefSeq database of bacterial genomes with RBD sequences in search of prophages with these motifs. We predicted the capsular tropism of the corresponding prophages by retrieving the CLT sequence of the host bacteria and inferring the capsular phenotype (Wyres et al., 2016). Then, we compared this prediction with the consensus CLT of the RBD cluster. We made enough predictions (> 30) for 18 of the 24 RBD clusters (79 %), which encompassed 15 distinct CLTs (KL2/13, KL3, KL22/37, KL23, KL25, KL30, KL35, KL47, KL57, KL63, KL64, KL102, KL106, KL126, and KL140). For the remaining clusters, we did not get enough hits with a confident CLT showing that these RBDs are rare in the prophages of typeable bacteria. Overall, we made ∼10.000 predictions, distributed in 3824 RefSeq bacterial genomes. The 18 RBD clusters had an average predictive power on CLT tropism of 35%, although this value varied widely, from 0 % to 100 % depending on the cluster. We found a statistically significant capsular tropism prediction for 13 clusters (72 %). Predictabilities > 98 % were obtained for four RBDs-CLT associations (KL2/13, KL30, KL57, and KL63; **Figure 4C**). Interestingly, it was possible to obtain accurate predictions (> 99 %) even beyond the *Klebsiella* genus (e.g. *Escherichia/Shigella*, for CLTs KL2/13, KL30, KL35, KL47, KL57, and KL63; 4/4, 90/91, 6/6, 43/43, 80/81 and 23/23 correct predictions, respectively).

For the five RBDs in which the predicted CLT based on prophages showed no significant association with the consensus CLT of the cluster analysis, we analyzed the distribution of hits to determine which factors might explain this failure. First, for RBD1_KL22/37, most hits (56.36 %) were assigned to KL25 instead of to KL22/37. Interestingly, capsule recombination between both CLTs has been previously identified (Haudiquet et al., 2021). Second, for RBD8_KL102, we found that 55 % of the predictions pointed to bacteria from the KL38. In this case, the RBD_KL102 representative sequence showed 35 % amino acid identity and 84% query coverage with the depolymerase of phage ϕKp34, which infects KL38 (Bonilla et al., 2021), revealing a potential cross-reactivity or an evolutionary association between the abilities to infect both CLTs. Next, clusters RB64 and RBD65 were both present in phages infecting KL23, but only prophages with identity to RBD65 were found in bacteria from KL23, suggesting that the latter was the active depolymerase. The same situation was observed in the RBD83_KL140. Finally, for the RBD87 cluster, the predicted CLT based on prophages was KL14 (72 %) instead of KL126 but the link between these two CLTs remains unknown.

### Post-entry restriction factors determine infection outcome

Spot tests provide a fast method for examining host tropism but they do not reveal productive infection (Domingo-Calap et al., 2020; Hyman and Abedon, 2010). To address this, for each of the 124 positive spot tests, we determined phage virulence by measuring optical density in liquid cultures. We detected a significant reduction in host density relative to non-inoculated controls in 94 of the 124 assays (76 %). Of the 30 negative pairs, five provided evidence of productive lytic infection by the progeny assay, and hence were also classified as virulent. Then, we reexamined host tropism by considering only the 99, out of 6319 (1.6 %), phage-host pairs yielding virulent infections in liquid culture (**Table S3**). Host breadth now ranged from 0 to 17 of the 138 tested strains. Again, the 42 non-S8/S9 phages were restricted to one or a few strains (median: 1, mean: 1.33), whereas S8-S9 phages exhibited a much broader range (median: 10.5, mean: 10.75). Host CLT was again the single factor that best predicted virulent infection, but with a considerably lower TPR than for spot tests (0.53 ± 0.01 for non-S8/S9 phages).

Therefore, spot tests seemed to be a better indicator of capsule-dependent host tropism than actual productive infection. To better understand this, we focused on the 25 phage-host combinations that yielded no-virulent infection despite the phages produced virulent infections in other strains (**Figure 5A**). As expected, non-S8/S9 phages induced capsule digestion in the 13/13 phage-host combinations in which the phage produced a spot, despite being avirulent (t-test: *P* ≤ 0.06; **Figure 5B**), showing the reliability of spot tests for assessing phage-induced depolymerization and capsule-dependent tropism. Interestingly, we observed successful irreversible adsorption for 16/25 combinations tested of which 14/19 included non-S8/S9 phages (**Figure 5B**).

**Figure 5.**
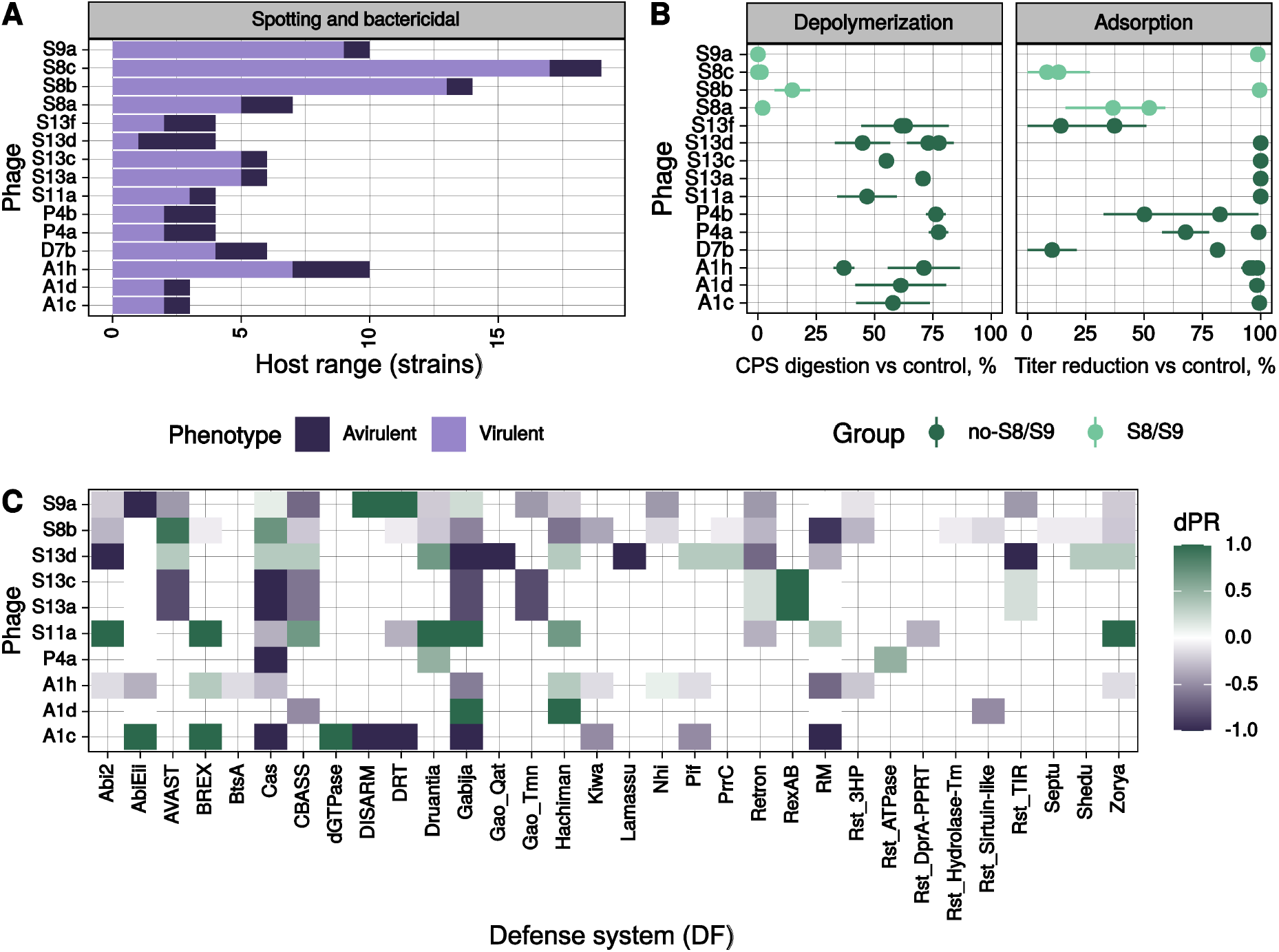
Post-entry restriction factors determine infection outcome. **A**. Summary of host ranges for phages that showed both virulent and avirulent infections. Virulent infections were considered when the phage was able to produce a spot and to reduce bacterial density or produce progeny. Avirulence was considered when despite spot formation, no productive infection and no significant effect on host density were observed. **B**. Quantitation of depolymerase activity and adsorption in 25 avirulent phage-host pairs. Depolymerase activity was quantified by comparing capsule polysaccharide levels in phage-treated versus untreated controls. Adsorption was quantified by measuring phage titer following inoculation relative to cell-free mock cultures (eclipse phase). Each point represents the mean +/-SEM of at least 3 replicates. **C**. Differential probability of resistance (dPR) for each phage-defense system. For avirulent host-phage combinations, only those in which adsorption was observed (panel B) were considered. Values of 1 indicate that the defense system was exclusively found in avirulent combinations. On the contrary, negative values indicate that the defense system was overrepresented in virulent combinations and thus probably did not contribute to the observed resistant phenotype.

To investigate the causes of avirulence despite efficient phage entry, we searched genomes for known anti-phage systems. We found no CRISPR spacers or resident prophages with significant identity to the adsorbed phages, suggesting no involvement of these forms of immunity. To perform a more systematic analysis, for each phage, we calculated the frequency of known bacterial defense systems in the different strains. We found 39 phage-defense system combinations which were more frequent in resistant (phage avirulence) than in susceptible strains (dPR > 0). Of them, 14 were exclusive of resistant bacteria (dPR = 1; **Figure 5C**). Overall, we found BREX, RexAB, Druantia, Hachiman, and dGTPase defense systems to be more frequently associated with resistant strains across phages (total dPR >= 1). Thus, even when phages stripped capsules and were adsorbed, there were intracellular mechanisms that protect bacteria. These should contribute to explain the lower sensitivity of CLTs for predicting the productive host range of phages. However, the mechanisms that govern these interactions, as well as the role of each defense system detected or unknown systems in the observed phenotypes, remain to be further characterized.

## DISCUSSION

Despite the relevance of phage-bacteria interactions, their study is limited by the low number of representative collections of bacteria and phages with a complete link between phenotypic and genotypic data (Lamy-Besnier et al., 2021). Here, we have performed a thorough analysis of the interactions between *K. pneumoniae* and its phages, using a larger phage-host interaction matrix than in previous studies (de Sousa et al., 2020; Townsend et al., 2021; Venturini et al., 2020). All the analyzed phages but one were novel and encompassed different families and undescribed taxonomical entities (Herridge et al., 2020), showing the importance of carrying out more extensive samplings to uncover the phage diversity (Göller et al., 2021; Kauffman et al., 2022; Maffei et al., 2021).

Given the tight relationship between phages and capsules, it is tempting to predict phage-bacteria interactions based on the CLT of the host and phage depolymerase sequences (Hryckowian et al., 2020; Sørensen et al., 2021). We found similar depolymerase domains between distant phages with overlapping CLT specificity, revealing a sizable diversity of depolymerases for each CLT. Effective switching of tail spike domains between phages has been shown under laboratory conditions (Latka et al., 2021) or concrete phages (Scholl et al., 2002). Our results suggest that tail spike domain swapping is frequent in nature, even across viral families. As a result, phage taxonomy was a poor predictor of host tropism, in contrast to some previous findings (Gencay et al., 2019; Sørensen et al., 2021) but agrees with recent work showing that phage host range is largely independent of phage taxonomy (Göller et al., 2021) or morphotype (Kauffman et al., 2022). Our results hint at the possibility of predicting phage capsular tropism only from depolymerase sequence information, a goal that we achieved partially and that should be further addressed in the future. Machine-learning approaches (Boeckaerts et al., 2021; Lood et al., 2021) in combination with deep mutational scanning studies to identify specificity regions in tail fibers (Huss et al., 2021) could help achieve this goal.

To test the ability of depolymerase RBD sequence analysis to predict capsular tropism, we used external data of prophage RBD sequences and the CLT of their hosts. It was thought that the depolymerase sequences of temperate phages exhibited limited identity to those of lytic phages (de Sousa et al., 2020), but our analysis reveals extensive sharing of depolymerase domains between both lifestyles. This may explain the high rate of recombination and suggests that temperate phages could be an important source of depolymerase domains for incoming lytic phages, as observed with other genes (de Sousa et al., 2021). We significantly predicted the capsular tropism for 13 CLTs, which represent around one-third of total CLT abundance in the global dataset of *K. pneumoniae*. The use of prophage RBD sequences to predict capsular tropism has several limitations. First, the host could undergo capsule swaps after prophage integration (Haudiquet et al., 2021), leading to false associations between the phage RBD sequence and the observed host CLT. Second, establishing actual RBD-CLT associations can be complex in prophages containing multiple RBDs. Third, some CLTs can be cross-reactive (Pieroni et al., 1994). Fourth, we should keep in mind that despite successful capsule binding, degradation, and adsorption, intracellular mechanisms can further restrict phage infection. This is in line with recent results in *Vibrio* sp., where phage infection varied across closely related strains due to post-adsorptive mechanisms (Hussain et al., 2021; Piel et al., 2021). Still, establishing solid associations between RBDs and pre-adsorptive phage activity might be useful from a phage therapy point of view, since degradation of the capsule by phages might help the immune system and/or antibiotics to clear infection (Fang et al., 2022; Gordillo Altamirano et al., 2021).

Despite current limitations, our results point to the feasibility of predicting phage infectivity in an encapsulated host of priority public-health concern such as *K. pneumoniae*. The results obtained here could be extended to other encapsulated pathogenic bacteria such as *Escherichia coli* and *Acinetobacter baumannii*, as their phages also contain depolymerase enzymes (Pires et al., 2016). Further work with *K. pneumoniae* is also needed because of the large unexplored capsular diversity, as revealed by the high number of depolymerases in phages with unassigned tropism. Some CLTs may be more abundant in environmental rather than in clinical strains and might be underrepresented in the databases. Also, different procedures could be used for phage isolation that might reveal different host range patterns. For instance, we expect that more broad-spectrum phages similar to those in the S8/S9 group could be discovered using acapsular *K. pneumoniae* for phage isolation. Further work will also be needed to better understand and predict how post-entry barriers to phage infection determine host tropism. More generally, comprehensive datasets with verified interactions similar to the data presented here are necessary to improve our understanding and predictability of phage-bacteria interactions in nature, and also for phage applications.

## MATERIAL & METHODS

### Phage isolation, purification, and sequencing

Environmental samples (n=9) were collected near a sewage water plant in Valencia (Spain) from soil and water. Isolation of candidate plaques was done as described previously (Domingo-Calap et al., 2020) using 36 clinical isolates of *K. pneumoniae* among which 19 had complete genome sequences (**Table S2**). Phages were amplified in 5 mL of LB broth for 3 h (150 rpm). After centrifugation (13,000 × *g*, 5 min), supernatants were passed through a 0.22 µm syringe filter to remove bacteria. Lysates were loaded into the upper reservoir of an Amicon filter device (100 KDa) and purified phages were recovered in 200 μL of SM buffer after two washing steps. Phages were ten-fold diluted in Turbo DNAse Buffer 1X (final volume of 100 μL) and 2 μL DNAse Turbo, benzonase, and micrococcal nuclease were added. The digestion was incubated for 1 h at 37°C and 1 μL of DNase Turbo was added during another hour. To inactivate DNases, 15 mM EDTA was used and incubated at 75°C for 10 min. To digest phage capsids, 10 μL of SDS 10% and 5 μL of proteinase K (20mg/mL) were added and incubated for 45 min at 55°C. DNA Clean & Concentrator 5-Kit (Zymo) was used to extract and purify the DNA. Sequencing libraries were prepared using the Illumina Nextera XT DNA kit and paired-end reads (2×250 bp) were generated in the Illumina MiSeq platform (MiSeq Reagent Kit v2). For transmission electron microscopy, a drop from purified phages was deposited onto a carbon-coated Formvar supported by a 300 mesh copper grid and air-dried for 30 min. Phages were negatively stained with 2% phosphotungstic acid and visualized under Jeol JEM-1010.

### Phage genome assembly and annotation

Raw reads were filtered with prinseq (Schmieder and Edwards, 2011) (-min_len 20 -min_qual_mean 20 -ns_max_n 30 -trim_qual_right 20 -derep 14) and assembled with Unicycler v0.4.7 (Wick et al., 2017). Alternatively, SPAdes v.3.14 (Bankevich et al., 2012) or MIRA4 (https://sourceforge.net/projects/mira-assembler/) were used with the apc script (https://github.com/tseemann/apc/blob/master/apc.pl) to check genome circularity. Pairwise average nucleotide identity (ANI) values were obtained using FastANI (Jain et al., 2018) (fraglen=50). After removing phage duplicates (ANI and alignment coverage, AC, >99%), intergenomic similarity (IGS) values were determined with VIRIDIC (Moraru et al., 2020). These were used to build a Neighbor-Joining tree with the ape package (Paradis and Schliep, 2019). Phage genomes were annotated with Prokka v1.14 (Seemann, 2014) using the database of Prokaryotic Virus Remote Homologous Groups (PHROGS) (Terzian et al., 2021) as suggested (http://millardlab.org/2021/11/21/phage-annotation-with-phrogs/). For visualization of phage genomes, phages were arbitrarily permuted at the small/large terminase subunit and genoplotR (Guy et al., 2010) was used to draw blastn comparisons (-evalue 1e-5). Temperate behavior was inferred by searching annotated lysogeny markers (integrases, excisionases, recombinase, transposase, transcriptional regulators/repressors and ParA/B genes).

### Phage classification

Phage genomes available at NCBI (PhageDB) were obtained with inphared (Cook et al., 2021) (10/02/2021, n=14037) and compared against isolated phages with FastANI (fraglen=500). PhageDB sequences with at least 70% ANI and 30% alignment coverage were kept for computing IGS values with our phages as above. The ANI.dendrogram function of the bactaxR package (Carroll et al., 2020) was used to define species and group clusters with IGS thresholds of 95% and 45%, respectively. ViPTreeGen (https://github.com/yosuken/ViPTreeGen) was used to compare phage genomes with all *Klebsiella* phages from the PhageDB.

### *K. pneumoniae* collection and genome analyses

The genome sequences of 1145 *K. pneumoniae* isolates (García-González N., et al., unpublished) were considered. Raw reads were trimmed with prinseq (Schmieder and Edwards, 2011) and Unicycler v.0.4.7 (Wick et al., 2017) was used for assembly. Isolates of each capsular locus type (CLT) were chosen based on assembly quality, confidence of the CLT serotyping, and the O antigen and ST diversity obtained with Kleborate (Wyres et al., 2016). Selected strains (160) were regrown and double colony-purified. To discard contamination, the newly purified isolates were typed by the *wzi* sequencing method (Brisse et al., 2013) and Sanger sequences were compared to the ones previously available from Illumina. This resulted in 138 *bona fide* strains including NTUH-K2044 (Wu et al., 2009) (**Table S2**). These were compared against representative sequences of the *K. pneumoniae* clonal complexes (de Sousa et al., 2020; Wyres et al., 2019). To do so, Snippy (https://github.com/tseemann/snippy) (--ctgs option) was used to obtain a core genome alignment of all sequences using NTUH-K2044 as reference. Positions with gaps in at least 10% of sequences were excluded with trimAl (Capella-Gutiérrez et al., 2009). IQ-TREE2 (Minh et al., 2020) was used to infer a core ML phylogeny under the best fitting model (Kalyaanamoorthy et al., 2017). PhageBoost (Sirén et al., 2021) was used to extract prophages. CRISPRCasTyper (Russel et al., 2020) was used to detect the presence of CRISPR systems. For the rest of bacteria defense systems, we used Defense-finder (Tesson et al., 2022) with default options. Finally, to determine the presence/absence of putative phage receptors, the genomes were annotated with Prokka v1.14 (Seemann, 2014). Proteins of phage receptors were obtained (Zhang et al., 2020) and a fasta file generated. This file as well as the 138 bacteria proteomes were used as input for PIRATE (Bayliss et al., 2019) to obtain a presence/absence matrix. Additionally, a semantic search was done in order to obtain divergent or additional outer membrane proteins (see Data availability).

### Phage host tropism

Firstly, phages were diluted in a 96 deep-well plate (>10^7^ PFU/mL) using a chess-board pattern. Spot tests were performed by adding 2 µL drops of each phage to the bacterial lawns of the 138 *K. pneumoniae* strains in 0.3 % top-agar LB media (Kauffman and Polz, 2018) and incubated at 37°C overnight. Three independent temporal blocks were performed. A case was considered as positive when at least two replicates showed a positive interaction (clear spot, turbid spot, or plaques). Every positive case was confirmed by the planktonic killing assay. Briefly, bacteria were streaked from glycerol stocks and grown to an exponential phase. Adjusted exponential cultures and phage stocks were used to inoculate 200 μL of LB in a 96-well plate, resulting in 107 CFU/mL and 108 PFU/mL, respectively. Plates were incubated at 37°C with shaking (60 rpm) in a Tecan microplate reader Infinite 200 and OD_600_ was recorded every 20’ for at least 16h. A positive bactericidal effect was assigned when a delay or change in bacterial growth was observed between the control and phage-treated bacteria in at least two replicates. Finally, for combinations in which a spot was observed but a bactericidal effect in liquid was not confirmed, phage progeny and adsorption were measured. At least 3 individual colonies were picked and grown in LB broth until they reached 107 CFU/mL. Bacteria were then infected with ∼105 PFU/mL and incubated for 3 hours at 37°C with orbital shaking (250 rpm). Then, plates were centrifuged (4000 x *g*, 10 min) and serial dilutions of supernatants were used to titrate phage progeny in the amplification strain of each phage by dotting assay. A positive progeny production was considered if phage exceeded 10-fold over free-cells control within 3 h. From these results, progeny production was calculated using the following expression: 100*(1-A/Ao), being A and Ao the phage PFU/mL with or without bacteria, respectively. For combinations where phage was not totally adsorbed after 3 h, the incubation time was shortened to 30 min to avoid interference with newly produced particles. The same procedure was used to determine irreversible adsorption but, after co-incubation of phage and bacteria, the mixture was vigorously vortexed and centrifuged (13,000 x *g*, 5 min). Positive adsorption was considered when the mean free-particle titer dropped > 80% compared to host-free controls.

### True positive rate (TPR) calculation

To predict the effect of bacterial and phage genetic traits on positive infections, the statistical sensitivity (i.e. TPR) was calculated. For bacteria, the infection matrix resulting from the spotting assay was used as input. Briefly, for each host trait (e.g. CLT of bacteria) and host trait status (e.g. KL2), a reference strain was selected and its pattern of infection determined. This infection profile was compared with the pattern of infection of other bacteria with the same trait status (e.g. KL2). The selection of the reference bacteria was repeated 100 times to avoid biases. The resulting contingency table of positive and negative infections was used to calculate the TPR as follows: TPR=P(S¦SR)/[P(S¦SR)+P(R¦SR)], being S|SR sensitivity to phages given sensitivity in the reference bacteria and R|SR resistance to phages given sensitivity in the reference bacteria. To determine the null TPR distribution of each trait, the reference strain was randomly selected within the whole dataset (e.g. any CLT). When the host trait represented binary data (e.g. presence/absence of a receptor), the TPR was estimated from the presence level. This process was repeated 3 times to estimate the standard error. For the TPR of phage traits, the CLT infection matrix was considered instead of individual phage-bacteria pairs. We considered that a phage was able to infect a CLT if at least one strain of the corresponding CLT was infected by the spotting assay or was referenced in the literature. Therefore, for each phage trait (family, group, or presence of a certain RBD), a reference phage was selected and the TPR calculated as above.

### Phage depolymerase activity

Aliquots of 2 mL of overnight bacterial cultures were treated with 10 μL of phage (MOI >=1) or 10 μL of SM buffer. After ON incubation at 37°C with moderate shaking (150 rpm), cultures were centrifuged (18,000 x *g*, 5 min) and washed twice with PBS. Then, the capsule was extracted as described previously (Domenico et al., 1989) with modifications. Washed cells in 0.5 mL of PBS were mixed with 100 μL of capsule extraction buffer (500 mM citric acid pH 2.0, 1% Zwittergent 3-10) and vortexed. The mix was heated at 56°C for 20 min and cellular debris was removed by centrifugation (18,000 x *g*, 5 min). To precipitate the capsule, the supernatant was mixed with ethanol to a final concentration of 80% (v/v) and incubated for 30 minutes at 4°C. The precipitates were collected by centrifugation (18,000 x *g*, 20 min, 4 °C), air-dried, and resuspended in 200 μL of water. CPS was incubated for 2 h at 56°C (Buffet et al., 2021) before quantification by the uronic acid method (Blumenkrantz and Asboe-Hansen, 1973). The capsule suspension was mixed with 1200 μL of borax solution (12.5 mM disodium tetraborate in H_2_SO_4_), kept on ice for 10 min, followed by incubation at 95°C for 10 min, and immediate cooling on ice for another 10 min. The absorbance of the sample at 520 nm was measured after the addition of 20 μL of 3-hydroxybiphenyl in 0.5% NaOH. A standard curve using glucuronic acid was used to calculate uronic acid concentrations for each quantification batch. Capsule reduction (%) was calculated by comparing the uronic acid content of phage-treated versus untreated bacteria. This procedure was repeated at least twice for each phage-bacteria combination tested. Phage depolymerization activity was calculated using the following expression: 100×(1-C/C_o_), being C and C_o_ the amount of bacterial glucuronic acid (μg/200μL) with or without phage, respectively. As positive controls of this assay, several phage-bacteria combinations in which capsule reduction was checked by microscopy were used (Domingo-Calap et al., 2020). The presence of haloes was checked by visual inspection of phage plaques after overnight incubation in at least one representative bacteria of a CLT.

### Search for depolymerases sequences

To enlarge our dataset of depolymerases analyzed, *Klebsiella* phages from the literature with known tropism were included (see Data availability). For those in which the proteins were not available at the NCBI, the phage genomes were assembled and annotated as described above. All phage proteins were compared against well-characterized (Pires et al., 2016; Latka et al., 2019) or experimentally validated depolymerase sequences (see Data availability) with blastp (e-value threshold of 10^−5^). Then, hhblits (PDB70) (Steinegger et al., 2019), InterProScan5 (Jones et al., 2014) and Phyre2 (Kelley et al., 2015) were used to verify the presence of the enzymatic domain. Alternatively, Pfam depolymerase domains previously available (Pires et al., 2016) and hits against signatures/profiles which contained the terms ‘depolymerase’, ‘pectin’, ‘pectate’, ‘sialidase’, ‘levanase’, ‘xylosidase’, ‘rhamnosidase’, ‘dextranase’, ‘alginate’, ‘hyaluronidase’, ‘hydrolase’, ‘lyase’, ‘lipase’ and ‘spike AND beta-helix|hydrolase’ in every CDS were tested. No additional depolymerases apart from those obtained with the blastp search were found. Proteins with less than 200 total residues and fewer than 100 aligned residues to an enzymatic domain (Latka et al., 2019) as well as hits of peptidoglycan enzymes/endolysins were removed. Tail fibers with seemingly enzymatic activity were therefore referred to as tail spikes.

### Delimitation of receptor-binding domains (RBDs)

To focus on regions driving host tropism, anchor domains of tail spikes were removed. These domains are conserved among related phages and do not confer specificity (Latka et al., 2019). To do so, tail spikes were compared against all phage proteins of the PhageDB and proteins of our phages with blastp (-evalue 1e-5 -max_hsps 1 -length 100). Hits were considered as intra-genus if the corresponding phages scored at least 70% ANI in 40% of their genome with FastANI (see above). Tail spikes of the same genus were used to build a sequence alignment with MAFFT (--adjustdirection). The conserved part of each protein alignment was obtained with Gblocks (-b3=4) (Castresana, 2000) and removed with subseq seqtk (https://github.com/lh3/seqtk). If more than one block was selected, the first block was considered as the anchor domain. If a conserved block could not be assigned or was < 20 aa, the whole protein was retained. For eight cases, the anchor domain was delimited after visual inspection of the alignment. The remaining length of the tail spike was referred to as the receptor-binding domain (RBD). To determine clusters of RBDs, all versus all blastp comparisons were performed (-evalue 1e-5 -max_hsps 1 -qcov > 40) and the bactaxR package used (Carroll et al., 2020) as above (identity threshold=50).

### Cross-validation of predicted tropisms using prophage sequence data

RBD clusters with capsular-tropism signal, that is, clusters where at least two-thirds of the sequences were in phages with shared CLT tropism, were extracted. Representative sequences of clusters were used to perform a search against the complete RefSeq protein database of bacteria with blastp (-evalue 1e-5 -max_hsps 1 -qcovs 0.4) to obtain putative prophage regions. Protein accessions of hits were used to retrieve the bacterial assembly accession and the corresponding taxonomy with efetch. Bacterial assemblies were downloaded from the NCBI ftp site and subjected to Kleborate (Wyres et al., 2016) to determine the CLT of each assembly. Bacterial assemblies with ‘None’ or ‘Low’ confidence were removed from the analyses. Then, the CLT assigned to each RBD from the cluster analysis was compared to the CLT of the bacteria with the residing prophage. To test for statistical significance, a random CLT was assigned to each bacterial assembly with probability equal to the frequency of each CLT in the *K. pneumoniae* RefSeq database (accessed: 27/04/2021). Then Fisher’s exact test was used to compare the proportion of correct predictions made in randomized versus actual data.

### Mechanisms of avirulent infections

The adsorbed phages were compared by blastn with bacteria residing prophages (e-value 1e-5, min identity 80%, and alignment length >= 99 nt) to determine possible superinfection exclusion. CRISPRCasTyper (Russel et al., 2020) was used to identify CRISPR arrays which were compared against phages with blastn-short (coverage and identity thresholds of 90% and e-value 1e-5) (de Sousa et al., 2020). The presence of each defense system (DF, n=42) was compared with the outcome of phage virulence (V) or avirulence (A) in that particular strain. Given this, the differential probability of resistance (dPR) for each phage-defense system was calculated as follows: dPR= P(DF|A) - P(DF|V), with values close to 1 indicating an overrepresentation of the DF in resistant strains (phage avirulence). On the contrary, negative values indicate that the DF was found overrepresented in susceptible strains (phage virulence) and they probably do not contribute to the observed resistant phenotype.

## Data and code availability

The FASTQ files of the 70 *Klebsiella* phages sequenced are deposited in the European Nucleotide Archive (*ENA*) under Bioproject PRJEB46367. The genome sequences of the 46 phages detailed in this study are deposited in NCBI GenBank according to the Accession numbers provided in Table S1. The sequences of the bacterial strains used are deposited in the ENA with Accession numbers provided in Table S2. The databases of depolymerases, phage receptors and *Klebsiella* phages from the literature as well as scripts used for True positive rate (TPR) calculations are available at https://data.mendeley.com/datasets/c696dvvynf/draft?a=88c9df17-13cf-408a-964c-52b19fedcbc6. Additional raw data or custom code used are available from Beatriz Beamud upon request.

## ACKNOWLEDGMENTS

We thank the Networked Laboratory for Surveillance of Antimicrobial Resistance (NLSAR) of Comunitat Valenciana, Ana Djukovic, Beatriz Herrera, Carles Úbeda and InfectERA Consortium for providing bacterial strains and access to genome sequences. We thank professor Jin Town Wang for the NTUH-K2044 strain. We thank David Saiz and Lourdes Tordera for help with strain collection and depolymerization assays, respectively, Concha Hueso, Nuria Jiménez and Loles Catalán for technical assistance and the Centro de Investigación Príncipe Felipe microscopy service for TEM. This research was funded by ERC grants 724519 - Vis-a-Vis and 101019724 - EVADER to R.S., project PID2020- 118602RB-I00 (Spanish MICINN), project BFU2017-89594R (Spanish MICINN) to F.G.C and project PROMETEO2016-122 (Generalitat Valenciana) to F.G.C and R.S, ESCMID Research Grant 20200063, project PID2020-112835RA-I00 funded by MCIN/AEI /10.13039/501100011033, and project SEJIGENT/2021/014 funded by Conselleria d’Innovació, Universitats, Ciència i Societat Digital (Generalitat Valenciana) to P.D-C. P.D-C. was financially supported by a Ramón y Cajal contract RYC2019-028015-I funded by MCIN/AEI/10.13039/501100011033, ESF Invest in your future. B.B was funded by a PhD fellowship from Spanish MCIU FPU16/02139.

## AUTHOR CONTRIBUTIONS

B.B. contributed to conceive the project, design research, perform the experiments/computational analyses, data analysis, and manuscript writing. N.G.G provided resources and helped with some computational analyses and visualization. M.G.O provided resources and helped with strain collection. F.G.C. contributed to design research, resources, manuscript revision, and supervision. P.D.C isolated the phages and contributed to conceive the project, design research, manuscript revision, and supervision. R.S. contributed to conceive the project, design research, data analysis, manuscript writing and revision, and supervision.

