## Supplementary Information for "Genetic determinants of host tropism in *Klebsiella* phages"

**SUPPLEMENTARY MATERIAL**

**Table S1. General features of the 46 novel bacteriophages**. Phage ID, phylogenetic classification in 13 groups, genome size and circularity, sequencing depth, GC %, CDS, putative lysogeny genes, taxonomic family, subfamily, and genus according to IGS values with the closest phage are provided. Additionally, formation of plaque haloes, susceptible CLTs and number of depolymerase domains are also shown. REP: transcription repressor/regulator, PAR: ParA/B genes, REC: recombinases. Taxonomy ranks marked with a star are new possible taxonomic ranks.

**Table S2. General features of the 138 *Klebsiella* clinical strains.** The strain, sequencing details, virulence and antibiotic resistance scores, *wzi* allele, CLT and OLT inference, number of prophages, CRISPR loci, and presence of defense systems and secondary receptors are provided. The presence of a capsule was determined by visual inspection of colonies using light microscopy.

**Table S3. Phage-bacteria positive combinations.** The strain used for phage isolation is indicated. Spotting: positive by the spot-assay. Virulent: positive by the killing planktonic or progeny assay.

**Table S4. Clustering of phage RBDs with putative depolymerase activity from this work and the literature.** RBDs are identified by the phage name followed by the phage protein in which the domain was detected, and the positions considered after the removal of the anchor domain.

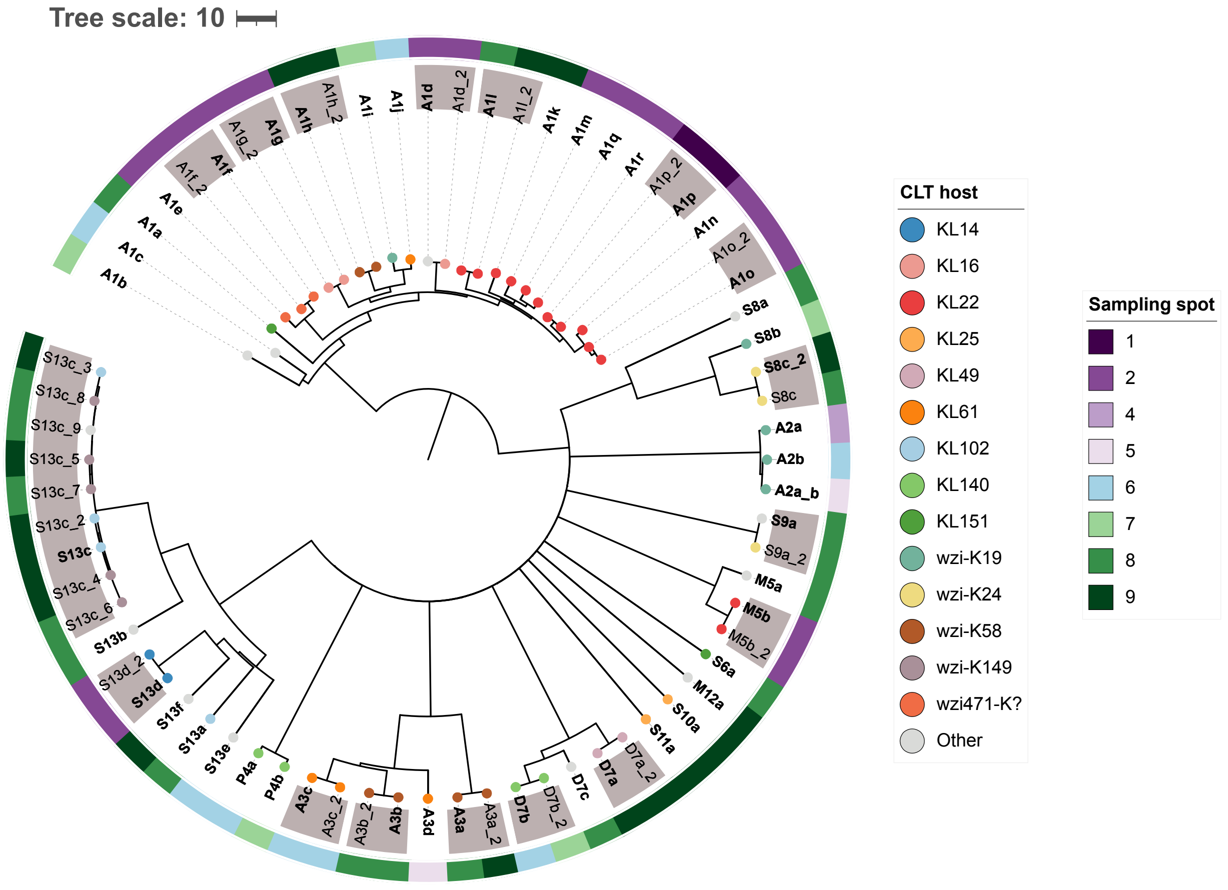


**Figure S1. Dendrogram of the 70 phage isolates.** A neighbor-joining tree was obtained from pairwise average nucleotide identity (ANI) values. Phages that were considered redundant are marked with gray boxes, and the selected representative of each phage strain is indicated in bold. This resulted in 46 distinct phages that were selected for further study. The capsular locus type (CLT) of the *Klebsiella* host used for phage isolation is indicated with colored circles at the tips of the dendrogram. CLTs with one occurrence are shown as 'Other'. ?: indicates *wzi* alleles whose correspondence with a CLT is unknown. The different sampling spots from which phages were isolated are indicated with colors outside the dendrogram. Tree scale denotes ANI distance (100-ANI).

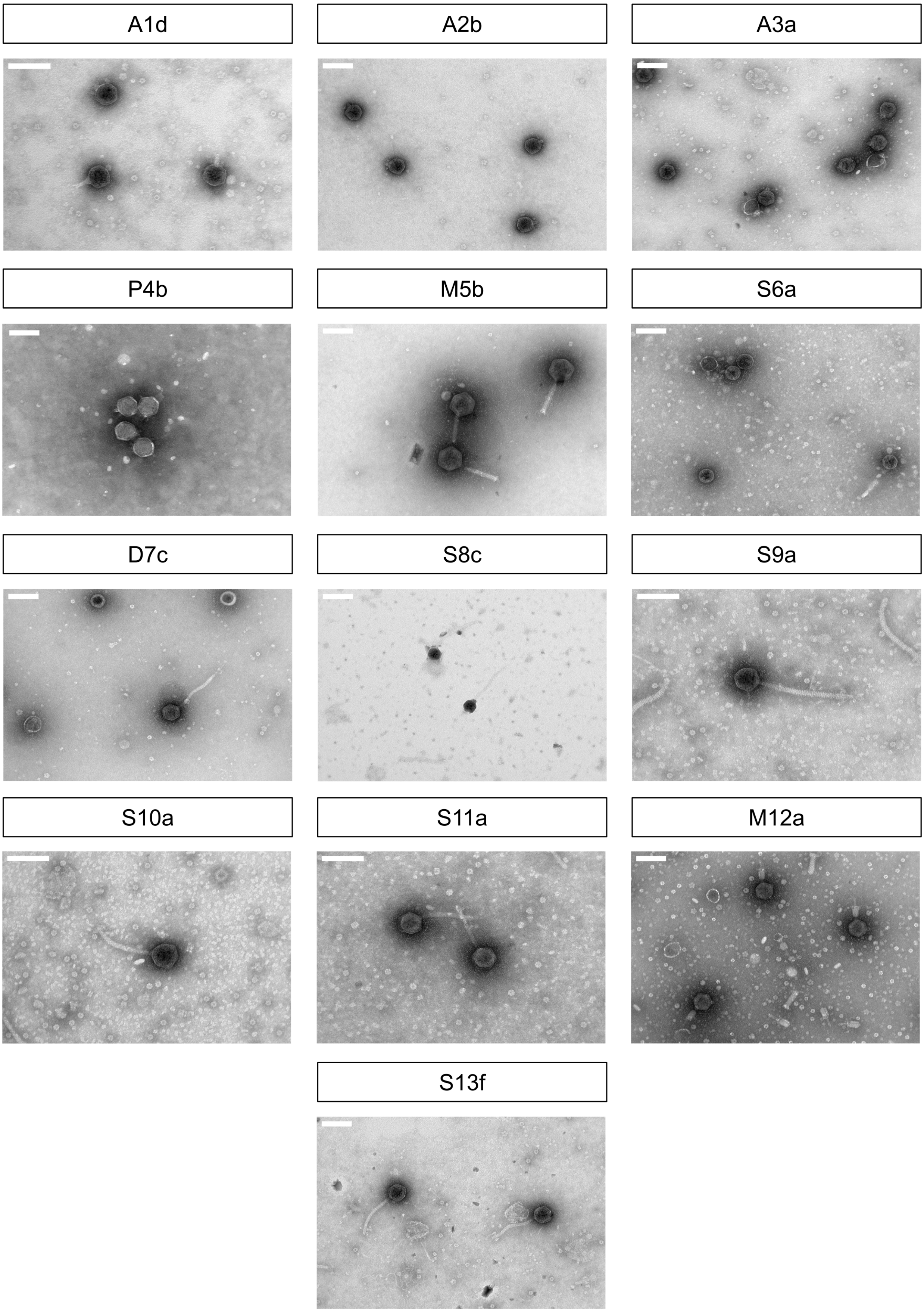


**Figure S2. Transmission electron micrographs of representative phages from each group.** Scale bar: 100 nm.

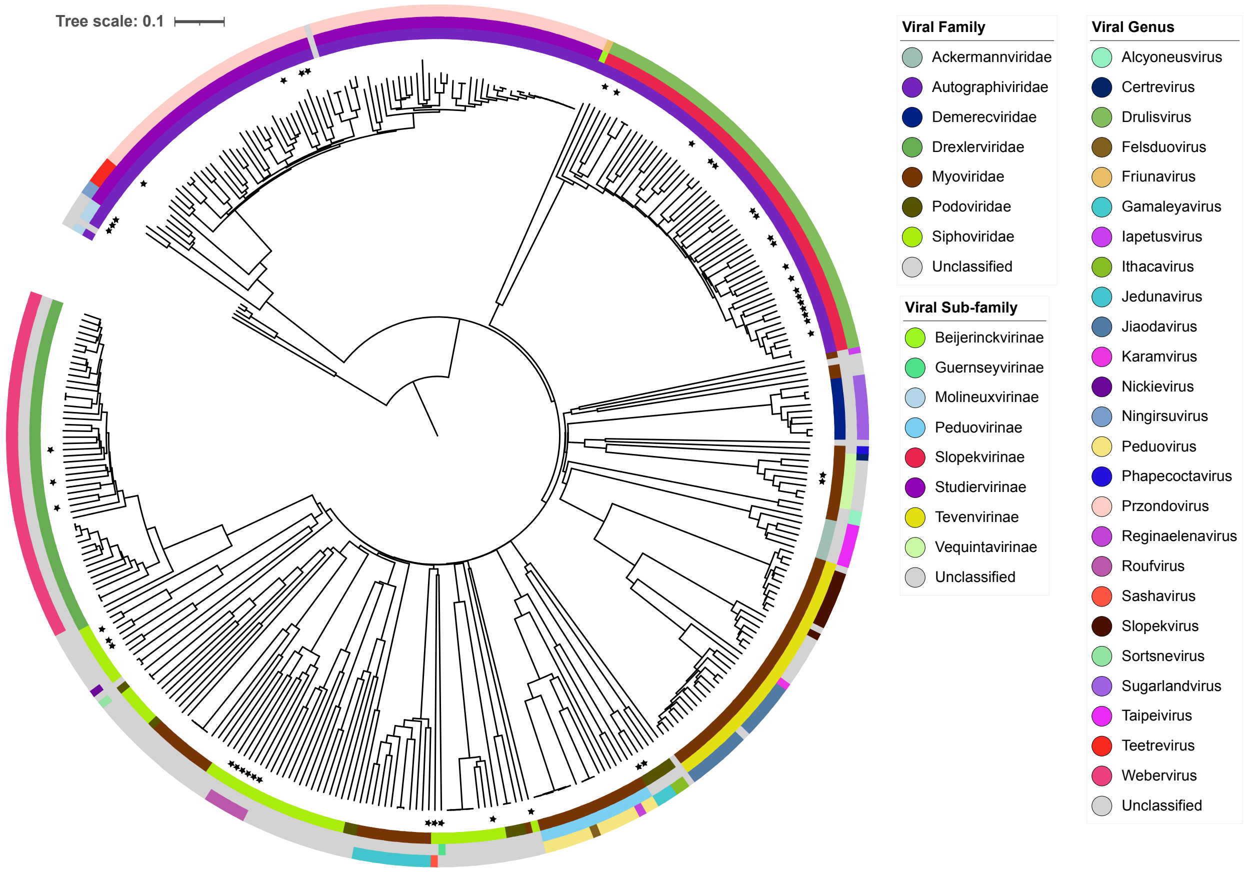


**Figure S3. Dendrogram of the new phages described in this study in the context of previously described *Klebsiella* phages.** Phage proteomic tree of all *Klebsiella* phages (February 2021), including the 46 newly sequenced phages (marked with a star) after computing genome-wide sequence similarities with tblastx. The tree scale shows global genomic distances. Clades of phages were colored based on available ICTV taxonomy.

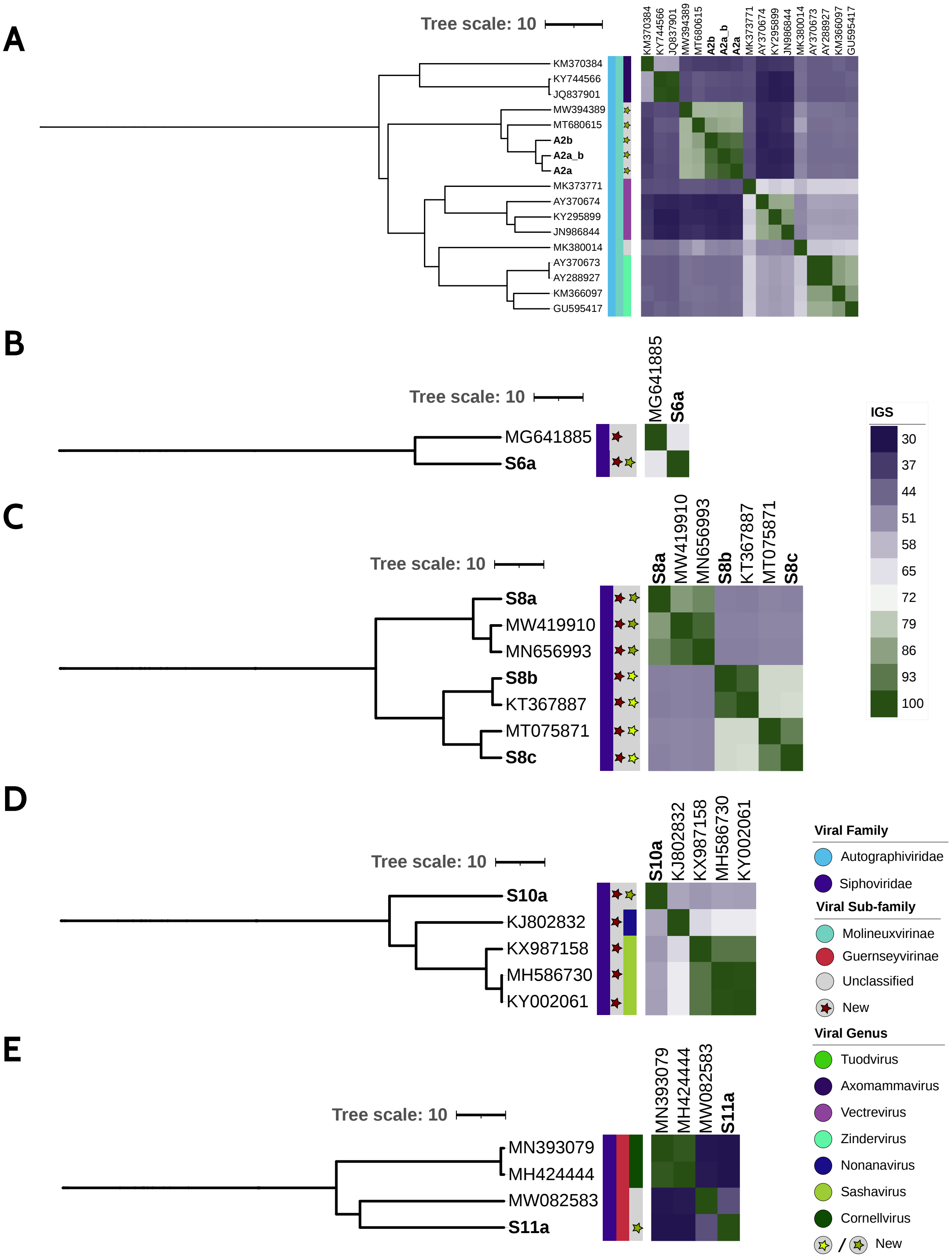


**Figure S4. Dendrograms of unclassified viruses based on IGS values.** Phages that were not classified at the subfamily and/or genus levels by similarity (IGS<45%) with any phage from the database were confronted against the NCBI (Accessed: 01/03/22) to exclude that new related phages had been deposited and classified since the initial analysis. Colored strips denote family, subfamily, and viral genus. New subfamily and genus suggestions are marked with a star and are based on updated ICTV thresholds (Turner et al. 2021). Phages from this work are in bold. **A.** Phages of group 2 (A2a, A2b, and A2a_b) represent a new genus, which includes *Proteus* phage PmP19 (MT680615) and *Klebsiella* phage vB_KpP_FBKp16 (MW394389), as their IGS values are >70%. **B.** Phage of group 6, S6a, is an orphan phage related at the subfamily level with *Salmonella* phage PMBT28 (MG641885) (IGS=65.3%). **C.** Phages of group 8 belong to the same subfamily (IGS > 45%) but to different genera: phage S8a belongs to the same genus as *Enterobacter* phages ATCEA85 (MN656993) and ATCEA23 (MW419910), whereas phages S8b and S8c are grouped with *Klebsiella* phages vB_Kp3 (same species as phage S8b) and vB_KleS-HSE3 (MT075871). **D.** Phage of group 10, S10a, is related at the subfamily level with *Sashavirus* and *Nonanavirus* phages, but represents a new genus. **E.** Phage of group 11, S11a, is an orphan phage within the *Guernseyvirinae* subfamily and a newly proposed genus. Tree scale denotes IGS distance (100-IGS).

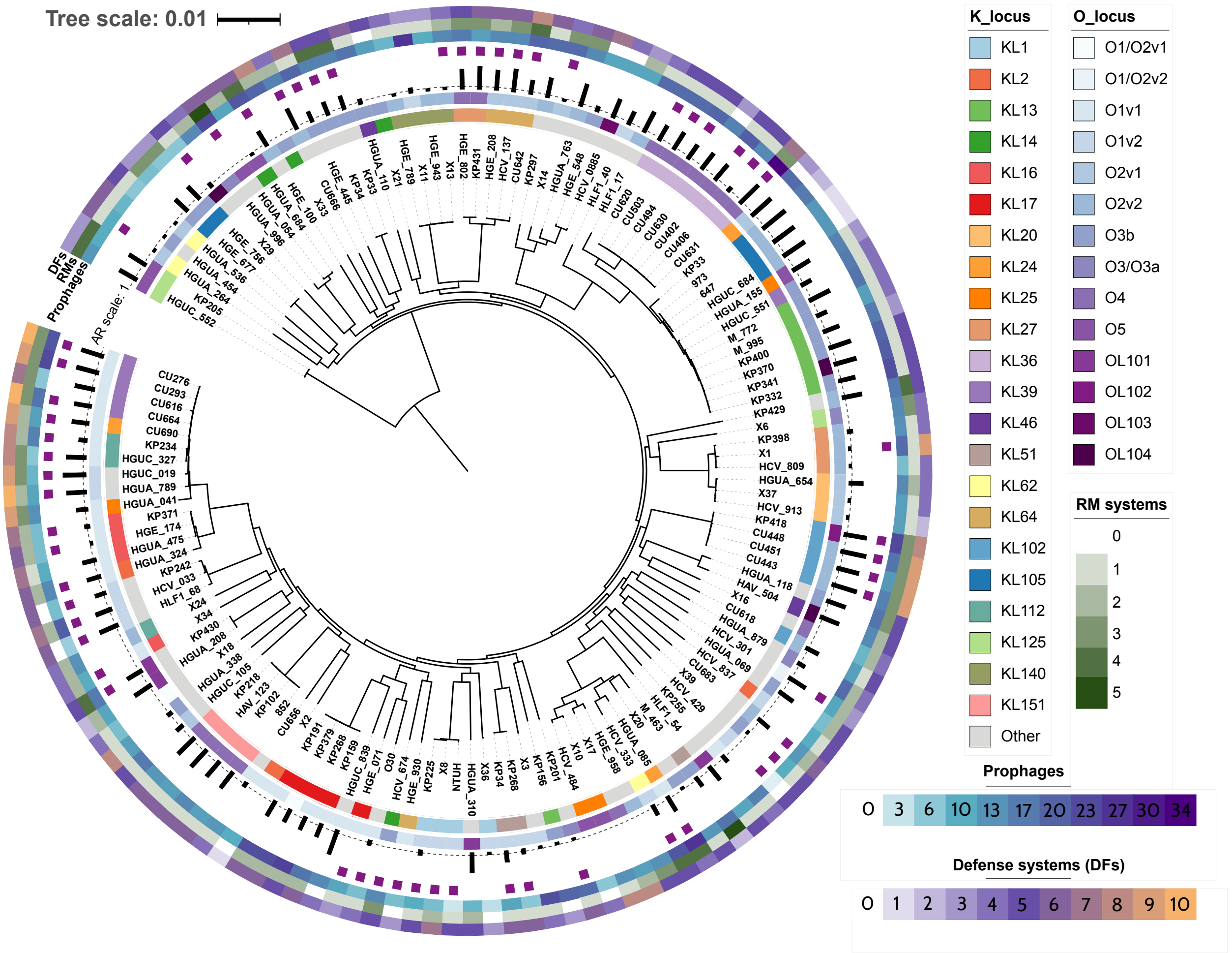


**Figure S5. Core genome maximum likelihood phylogeny of *K. pneumoniae* strains used in this study.** The innermost ring shows K-antigen prediction (capsular locus type, CLT) followed by O-antigen prediction (OLT). CLTs with fewer than three strains are shown as 'Other'. Black bars indicate the number of resistance genes to different antibiotic classes. Purple squares indicate the presence of CRISPR-Cas loci. The number of prophages, Restriction-Modification systems (RMs) and other defense systems (DFs) are also shown for each strain. Five strains that did not belong to *K. pneumoniae sensu stricto* (KP42, X22, CU371, KP205, HGUC_552) were omitted from this view.

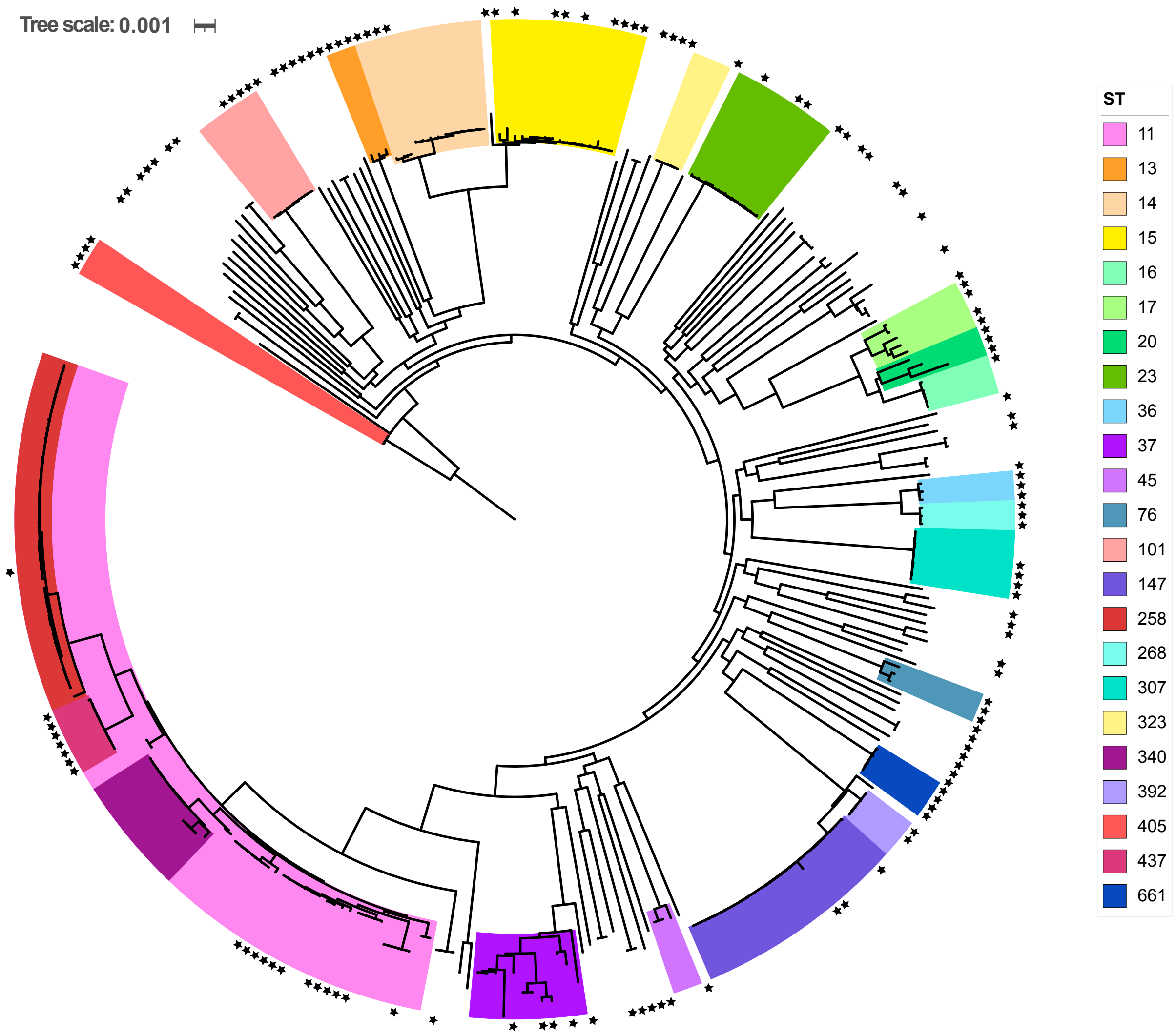


**Figure S6. Phylogenetic context of the *K. pneumoniae* strains used in this study**. A maximum-likelihood tree was constructed using the 138 strains included in this study (indicated with asterisks) and 172 representative sequences of each *K. pneumoniae* clonal complex. Colors represent major sequence types (STs). The five strains that did not belong to *K. pneumoniae sensu stricto* were excluded from this view.
